# Evidence for natural selection shaping the evolution of collective behavior among global *Caenorhabditis elegans* populations

**DOI:** 10.1101/2025.04.29.651269

**Authors:** Youn Jae Kang, Antonio Carlos Costa, Ryan Greenway, Priyanka Shrestha, Assaf Pertzelan, André EX Brown, Siyu Serena Ding

## Abstract

Animal behavior can diverge in natural populations in response to different environmental conditions, but if and how natural selection also shapes the evolution of collective behavior in groups of animals remains underexplored. With their cosmopolitan distribution and known collective behaviours, wild populations of *Caenorhabditis elegans* provide a powerful system to address how collective behavior could evolve across natural habitats on a global scale. We screened a panel of 196 genetically diverse *C. elegans* strains sampled from around the world, conducting aggregation behavior experiments and analysis to quantify natural variation among these populations. We found substantial variation in the spatial magnitude and the temporal dynamics of aggregation across strains, which were significantly explained by the elevation of the source habitats. Accounting for neutral evolutionary processes, our maximum likelihood population effects (MLPE) models further support a role of selection on aggregation. Furthermore, the two behavioral traits are highly heritable, and genome-wide association studies (GWAS) revealed a quantitative trait locus (QTL) containing several candidate genes associated with oxygen response and foraging behaviors. Our results showcase *C. elegans* aggregation as a collective behavior that has diverged globally across elevational gradients, and support that natural selection has shaped the evolution of this collective behavior.

## Introduction

Collective behavior is pervasive in the animal kingdom, but how collective behavior of animal groups evolves in nature remains elusive. Collective behavior arises from the interactions among multiple individuals in a group; this intrinsic complexity has traditionally been difficult to capture and quantify. However, rapid developments in behavior recordings and analytical techniques in recent years are increasingly facilitating rigorous quantification of complex behaviors (1–6). Behavioral data thus systematically captured and measured have enabled functional and mechanistic dissections of collective behaviors in several animal taxa (7–17) Recently, an increasing number of comparative studies reported natural variation in collective behaviors among different populations and species (17–27); however, few studies have investigated the mechanisms driving the evolution of collective behaviors among populations. Investigations of individual level behaviors have demonstrated that natural selection can drive the divergence of various animal behaviors among natural populations in response to different environmental conditions (28–35) but work on acorn ants and sticklebacks suggest that individual and collective behavior may have different evolutionary patterns (17, 18, 36) Does collective behavior also evolve under natural selection? Here we study how collective behavior may evolve across natural habitats on a global scale and place patterns of phenotypic variation in an environmental and genetic context using the nematode *Caenorhabditis elegans*.

*C. elegans* is a free-living bacterivore nematode species commonly found in decomposing organic matter (37). Recent genomic analysis of wild strains isolated worldwide suggests that this species has rapidly spread from its ancestral environments in the Hawaiian Pacific region out to the rest of the world in the past 100-200 years (38, 39), thus raising the question of how *C. elegans* behavior might have diverged in different environments around the globe. Neutral evolutionary processes, such as genetic drift and gene flow, would be expected to strongly influence the evolution of *C. elegans* traits given the species’ recent colonization history and predominately selfing reproductive strategy (38–41), but does natural selection also shape the evolution of collective behavior in this cosmopolitan species? The boom-and-bust life cycle of *C. elegans* in nature (42) implies ecological relevance of studying collective behavior in this species, as the animals experience rapid population expansion on food sources where thousands of individuals could be found in the same resource patch (43–46) subsequent resource exhaustion leads to population bust and dispersal to locate new resources, a process that can also occur collectively (47, 48) Here, we focus on the natural variation of *C. elegans* aggregation behavior on food to understand the evolution of collective behavior across global environments in this species.

Worms feeding on a bacterial food patch together can aggregate into clusters. Past work has examined this behavior in *C. elegans* and identified that variation in the *npr-1* gene underlies solitary versus gregarious phenotypes in lab-domesticated strains (44). A subsequent body of work has focused on understanding the molecular and cellular mechanisms (49–51), the algorithmic emergence (9), and the potential fitness implication (52) of aggregation behavior by contrasting the canonically solitary lab reference strain N2 with its gregarious counterpart, the *npr-1* mutant. However, behavioral variation amongst wild-collected strains has not been well-characterized. Meanwhile, a study of wild populations of another nematode species, *Pristionchus pacificus*, reveals a behavioral dichotomy between aggregation and solitary behavior across a regional elevation threshold (53), further prompting a detailed investigation of aggregation behavior among wild *C. elegans* populations. We hypothesize that aggregation has evolved in *C. elegans* across different habitats around the world, and that we can identify environmental and genetic correlates associated with this behavioural variation.

We conducted controlled behavior experiments in the lab to characterize aggregation behavior, first for the two canonical laboratory strains, and then for an extended panel of 196 genetically diverse wild strains isolated from around the globe. We developed sensitive quantitative representations for the spatial magnitude and the temporal dynamics of aggregation behavior, which revealed ample natural variation across populations. Accounting for phylogenetic structure among strains, we assessed the association between the behavioral traits and environmental factors, and found habitat elevation to be a strong predictor of both the magnitude and the dynamics of aggregation. We further partitioned out the expected role of neutral evolution using maximum likelihood population effects (MLPE) models to confirm an effect of selection on aggregation dynamics. Moreover, both behavioral traits are highly heritable, and genome-wide association studies (GWAS) reveal a quantitative trait locus (QTL) for aggregation dynamics that contains genes with known function in oxygen response and foraging behavior in *C. elegans*. Taken together, our results reveal natural variation in aggregation *C. elegans* populations and identify its environmental and genetic correlates, suggesting that this collective behavior has evolved in response to natural selection.

## Results

### Behavior assay and quantification effectively characterize aggregation behavior

We designed an aggregation behavior assay to experimentally capture variation in the collective behavior among different *C. elegans* strains: the two canonical laboratory strains N2 and *npr-1*, and one wild strain CB4856. We used forty age-matched individuals per assay to reduce inter-individual variation within the experiment and conducted 45-minute recordings on agar plates (∅ 35 mm) seeded with an OP50 bacterial patch (Fig 1a).

**Figure 1:**
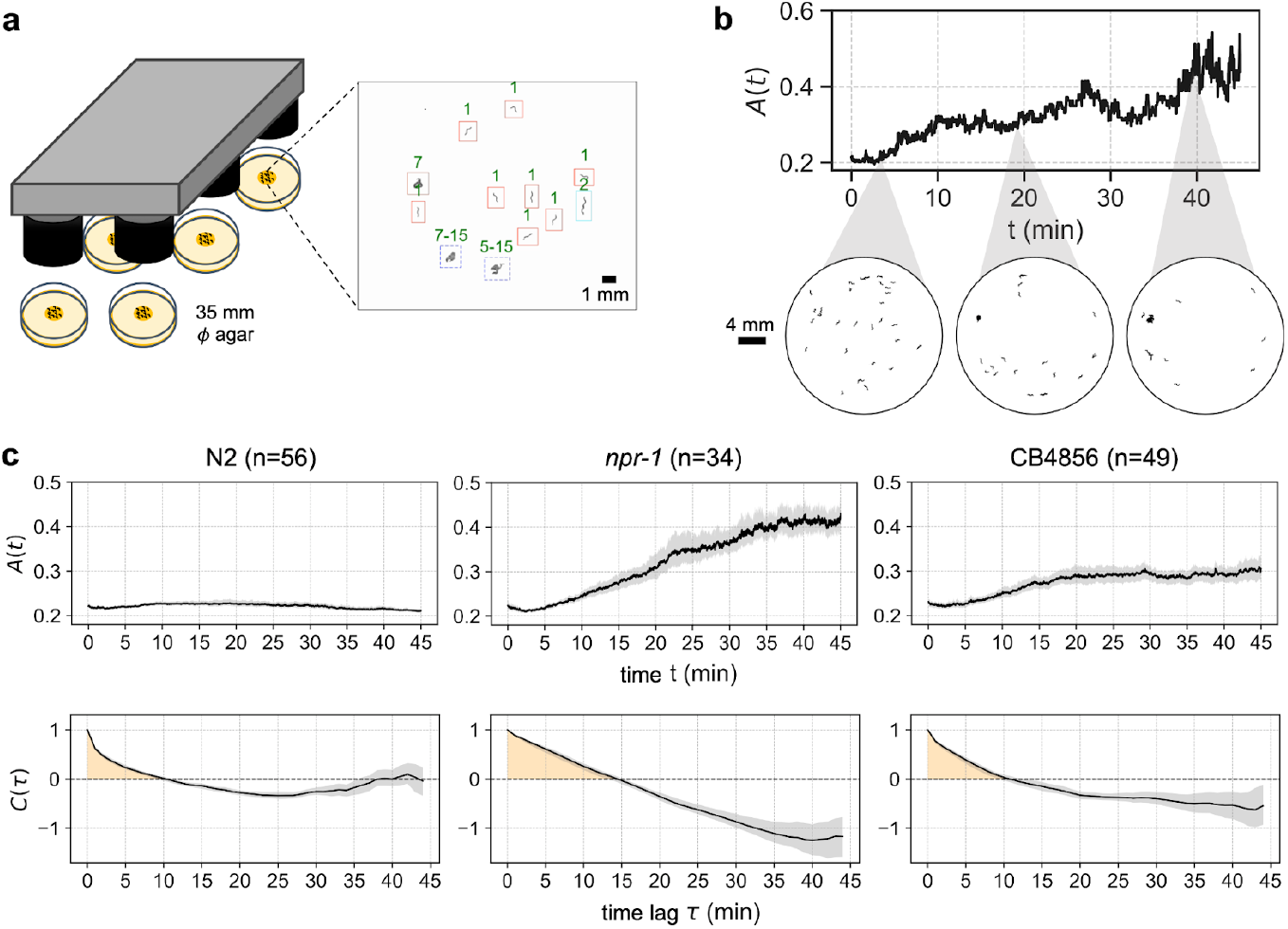
*C. elegans* aggregation behavior assay and quantification. a. Behavior assay to capture variation in aggregation behavior. The center of an agar plate was seeded with 75 µL of OP50 bacteria to form a circular patch. Forty worms were placed in the arena and recorded for 45 minutes using a multi-camera array. The image on the right depicts cluster size estimates from a sample frame. b. Sample aggregation time series from a recording of the *npr-1* strain. Top panel shows the magnitude of aggregation, defined as the inverse spatial entropy *A*(*t*); bottom panel shows snap shots of worm spatial distribution at given time points. c. Top panels show aggregation time series of the three representative strains: N2 is the solitary lab reference strain, *npr-1* is the gregarious knock-out mutant strain, and CB4856 is a gregarious wild strain. Bottom panels show the autocorrelation function of the corresponding *A*(*t*), to reveal differences in the temporal dynamics of aggregation behavior among the three strains. Yellow shadings depict the areas quantified as temporal persistence τ_A_ CB4856 exhibits intermediate characteristics in both aggregation magnitude and dynamics compared to N2 and *npr-1*

For behavior analysis, discrete object ‘blobs’ were segmented from the background using an adaptive local threshold. Single worms and multi-worm clusters were distinguished through a graph-based tracking algorithm that connected detected objects across consecutive frames and integrated morphological features with trajectory characteristics (see Methods). Cluster sizes were estimated using an iterative propagation method that exploited conservation constraints at splitting and merging events (see Methods). The resulting cluster size distributions from every frame were used to quantify the state of aggregation on a continuous scale over time.

Previous aggregation metrics such as the percentage of animals inside groups (44, 49–53) fail to capture spatial and temporal details to reveal natural variation amongst selected wild strains used in those studies; a recent improvement has been made for time-independent spatial distribution of individuals (9), but still ignores the temporal aspect of this dynamic collective behavior. To faithfully capture the fine behavioral variation and rich dynamics, we defined an aggregation metric that can be calculated on a per-frame basis: the inverse spatial entropy of the distribution of cluster size, *A*(*t*). If worms are tightly aggregated, the spatial distribution of worms deviates greatly from uniform distribution, hence the total entropy decreases and *A*(*t*) is high (Fig 1b). For the dynamics of aggregation behavior, we assessed the autocorrelation function &(τ) of the aggregation time series. If aggregation persisted more stably over time, the period and the timescale of the behavior was longer and the decay of the autocorrelation function was slower (Fig 1c).

We first assessed our behavioral metrics on the two laboratory strains with well-characterized aggregation behavior: N2 is the lab-domesticated reference strain known for its solitary behavior on food whereas *npr-1* mutants exhibit strong aggregation under the same conditions (44, 49–52). Indeed, *A*(*t*) remains very low throughout the recording duration for N2, with a short behavior timescale, while the *npr-1* mutant strain shows a drastic increase in aggregation magnitude over time and a longer behavior timescale (Fig 1c). We compared these results to a wild strain CB4856, commonly reported to be gregarious (44, 52). As expected, CB4856 worms show a high aggregation magnitude and a long behavior timescale, but with reduced values compared to the *npr-1* mutants (Fig 1c). Our quantification thus successfully captures the complex dynamics of *C. elegans* collective behavior in both domesticated and wild strains, and reveals fine differences between the two gregarious strains.

### Behavior quantification reveals natural variation in the aggregation behavior among wild populations

We hypothesized that aggregation has evolved across global habitats, and therefore predicted to find natural variation in the behavior. To widely sample the collective behavior of wild *C. elegans* populations, we applied the assay developed above to a set of 196 strains from *Caenorhabditis* Natural Diversity Resource (CaeNDR). The strains were collected from various natural habitats around the globe (Fig 2a) and are genetically diverse (54). The highest sampling efforts were in Europe (125 strains) and North America (34 strains), followed by Oceania (15 strains), Africa (9 strains), South America (5 strains) and Asia (3 strains), with the remaining 5 strains missing GPS information from the database (see Methods).

**Figure 2:**
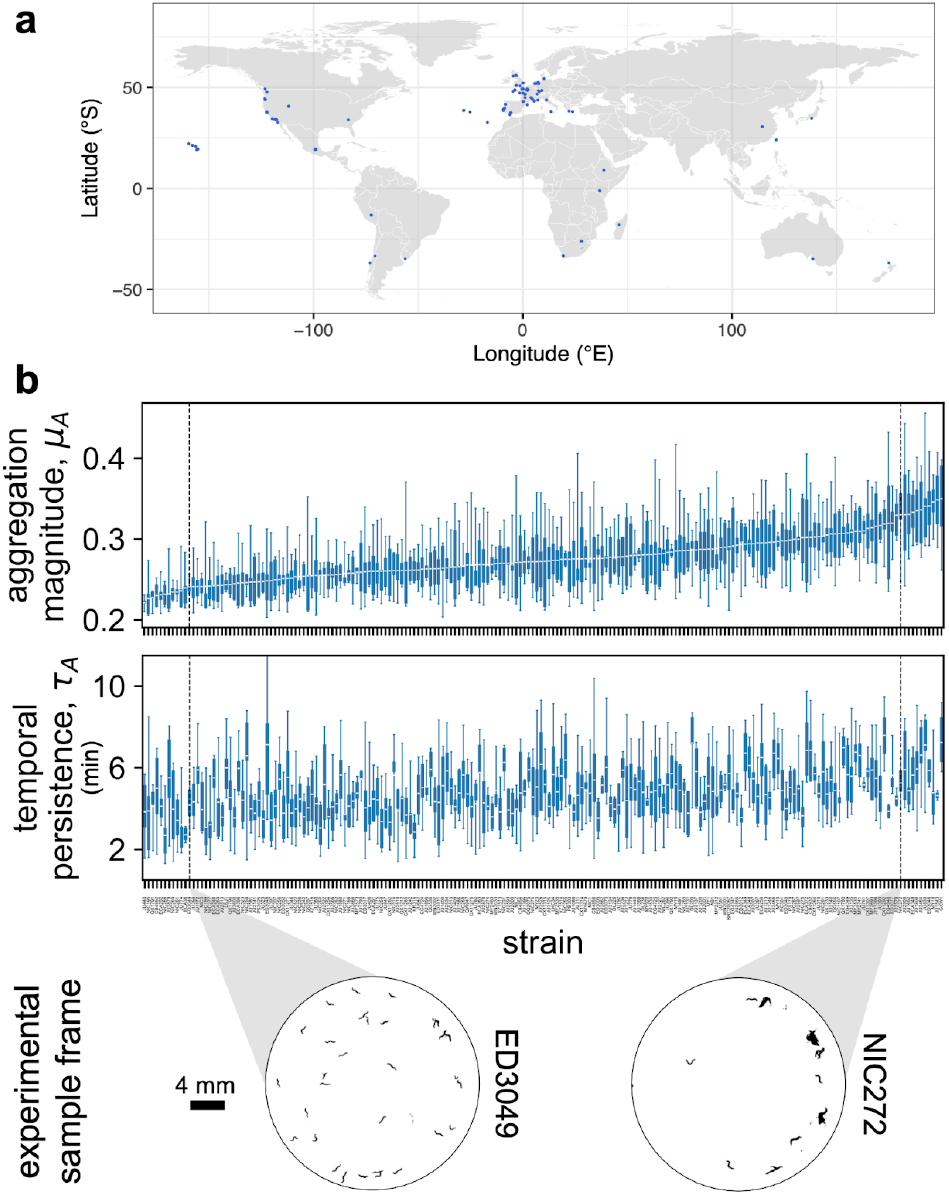
Natural variation in aggregation phenotypes. a. Geographical location of the isolation sites of the 196 wild *C elegans* populations. b. Aggregation metrics. Top panel shows time averaged aggregation magnitude *μ*_*A*_ quantified for each strain, organized in ascending order of the strain mean value. Middle panel shows temporal persistence τ_*A*_ quantified for each strain, organized in the corresponding order to the top panel. The boxplots display the mean and the interquantile range with minima and maxima of the experimental replicates per strain. Sample size of each strain can be found in Fig S2 and Fig S3. Bottom panel depicts sample aggregation behavior of strains on the lower and higher end of the population range for aggregation magnitude. A representative frame was chosen at around 40 minutes into the recording for both strains.

We found substantial phenotypic variation across the global panel of 196 wild strains in both aggregation magnitude and dynamics (Fig S2, Fig S3). To summarize the aggregation time series and the autocorrelation function of each strain, we extracted one scalar value from each function to obtain time averaged magnitude *μ*_A_ and temporal persistence τ_A_ for every experiment per strain (see Methods). Higher mean magnitude signifies a strain with more clustered behavior, and higher temporal persistence indicates a strain with higher behavior stability over time. We confirmed natural variation in both aggregation magnitude and temporal persistence in our global strain set (Fig 2b). Our two metrics thus effectively compress high dimensional behavior data into two concise trait values whilst maintaining the sensitivity to capture natural variation in aggregation, and support that this collective behavior has diverged among wild *C. elegans* strains.

### Variation in the aggregation traits is explained by elevational gradient

What is the evolutionary mechanism behind the observed global divergence in *C. elegans* aggregation behavior? To test whether environmental factors could be responsible for driving the evolution of this collective behavior via natural selection, we assessed the association between the aggregation traits and four categories of environmental variables: elevation, near-surface temperature, near-surface relative humidity, and moisture of the soil upper column (Fig 3a, S1). Ambient oxygen levels are known to affect aggregation behavior in domesticated *C. elegans* and in wild *P. pacificus*, where worms with a preference for low oxygen cluster more when ambient oxygen is high (49, 50, 53, 55, 56). Therefore we hypothesize that elevation, negatively correlated with atmospheric oxygen partial pressures (53, 57, 58), may also predict aggregation behavior in wild *C. elegans* populations. We chose the three additional environmental variables to reflect general characteristics of near-surface terrestrial habitats for *C. elegans*, in line with previous work that sought to identify correlations between genetic and climate variations in wild populations (55). Mean and standard deviation of the latter three variables were extracted over a 15-year period to represent the average and the fluctuation levels respectively (Fig 3a, S1), giving a total of seven environmental predictors for subsequent environment-phenotype association analysis (Fig 3a).

**Figure 3:**
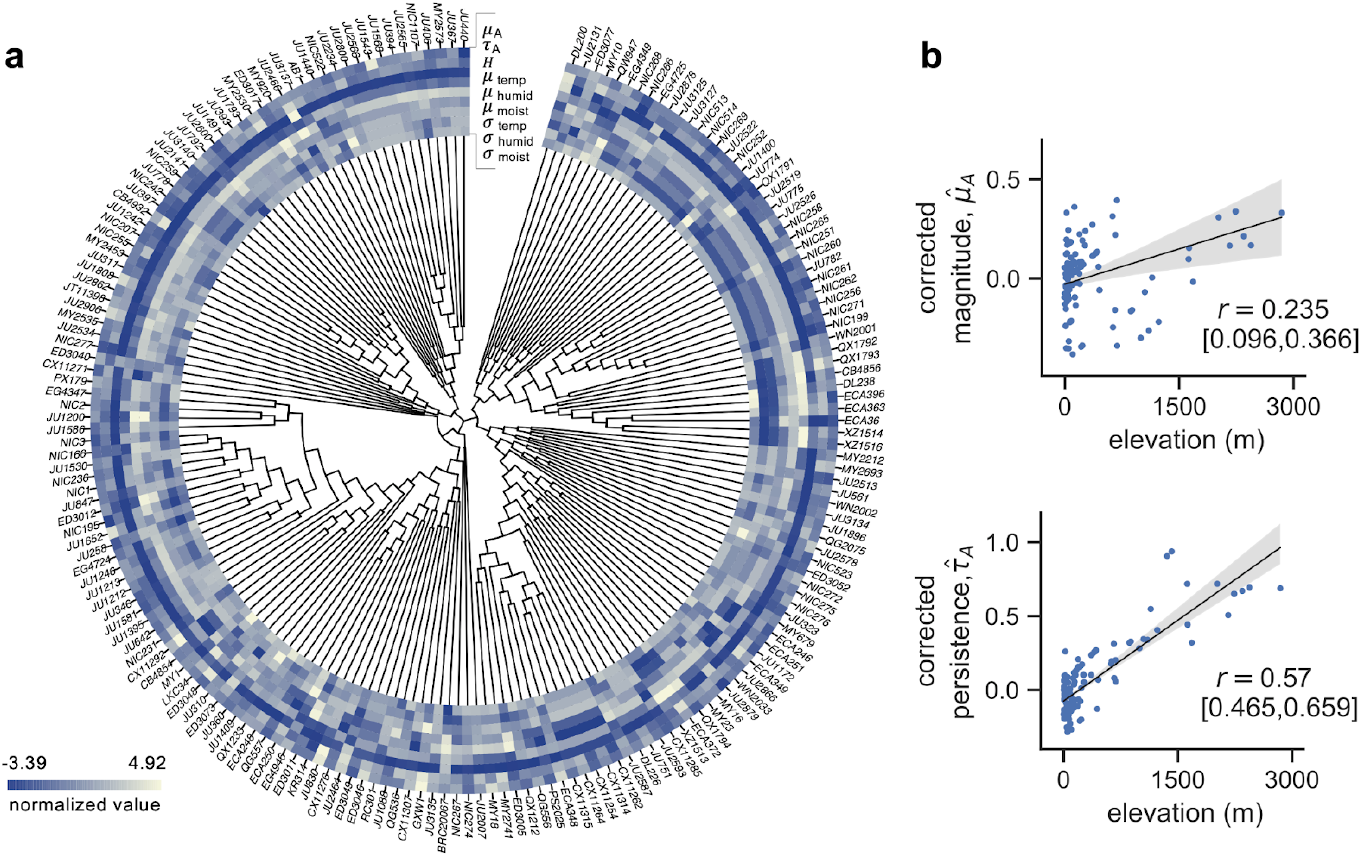
Environmental correlates of the natural variation in aggregation behavior. a. Phylogenetic tree and heatmap of the two aggregation traits (*μ*_*A*_, τ_*A*_) and the seven environmental predictors: elevation (*H*) and the mean and standard deviation of near-surface temperature (*μ*_*temp*_, *σ*_*temp*_), near-surface humidity (*μ*_*humid*_, *σ*_*humid*_) and upper column soil moisture (*μ*_*moist*_, *σ*_*moist*_). Each variable is z-score normalized. Association between the aggregation traits and the environmental variables was computed using the phylogenetic GLS model to account for the genetic structure among the wild strains. b. PGLS reports a significant effect of elevation on both aggregation traits. Top plot shows the correlation between elevation and mean aggregation magnitude; bottom plot shows the correlation between elevation and temporal persistence. Shaded areas depict 95% confidence intervals of the regression lines.

Our extended strain panel may contain hierarchical structure based on the phylogenetic relatedness among strains, leading to correlations between phylogeny and phenotype (59). Correcting for such hierarchical genetic structure, we used a Phylogenetic Generalised Least Squares (PGLS) model to assess the associations between the behavior traits and the environmental predictors: *μ*_*A*_ or τ_*A*_ is the response variable, (*H*_*elevation*_, *μ*_*temp*_, *μ*_*humid*_, *μ*_*moist*_, *σ*_*temp*_, *σ*_*humid*_, *σ*_*moist*_ are the predictors, and phylogeny is considered as a random effect (see Methods). Phylogenetically corrected behavior phenotype was computed as: 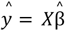, which assumes 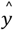 as the projection onto a space of environmental predictors in the absence of genetic random effects. Model selection was also performed to determine a set of environmental predictors that can best explain the behavioral variation. Akaike Information Criterion (AIC) was used as the score of model fit to compare between the models with different subsets of environmental predictors.

PGLS reported a positive and significant relationship between elevation and aggregation magnitude (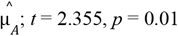; Table S2; Spearman’s *r* = 0.235 (CI: [0.096, 0.366]); Fig 3b top), and between elevation and temporal persistence (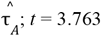; Table S4; Spearman’s *r* = 0.57 (CI: [0.465, 0.659]); Fig 3b bottom). This indicates that the strains from higher elevations exhibit higher aggregation magnitude and longer temporal persistence in our behavior assay. Interestingly, the best fit models of both aggregation traits frequently included the standard deviation of temperature as an environmental predictor alongside elevation, even though no significant effect was reported in PGLS for *σ*_temp_ alone (Table S3, Table S5). We assessed multicollinearity among the predictors and found that elevation was negatively correlated with the standard deviation of temperature (Pearson’s -0.487) (Table S6), indicating that as the elevation of the habitat increases, temperature fluctuations would generally decrease. Our results show that natural variation in the aggregation traits of our global strain set are significantly explained by a positive association with elevation and possibly by a negative association with temperature fluctuations in their natural habitats. The identification of a strong environmental predictor for aggregation suggests that the behavior may be under natural selection in wild *C. elegans* populations.

Due to the recent global spread of *C. elegans* and strong selective sweeps detected across much of the genome, genetic drift (due to their predominately selfing reproductive mode and founder effects) and gene flow (through human mediated dispersal) play a major role in shaping the evolution of this species (38, 39). As a means to disentangle the relative role of natural selection and neutral evolution (genetic drift and gene flow) in shaping the evolution of collective behaviors among populations, we used MLPE models to explicitly test the effects of genetics, geography, and elevation on behavioral divergence among sampled strains. If divergence in collective behaviors originates due to natural selection from elevational differences between sites, pairwise behavioral distances between strains should be correlated with pairwise environmental differences between collection sites (60, 61) Divergence in collective behavior stemming from neutral evolution should be evident from correlations between pairwise genetic and behavioral differences, so that more genetically similar strains are also behaviorally more similar, provided that genetic distances are not strongly influenced by environmental differences between sites (60, 61) We first confirmed that pairwise genetic distance between strains was best explained by the geographic distance between sampling locations (β = 0.02 ± 0.00, t = 38.76, p < 0.001; Table S8), and was not associated with differences in elevation between sites, indicating that geographically closer populations are genetically more similar, and that gene flow is not inhibited between populations from different elevations. We then tested for correlations between each aggregation trait and neutral genetic variation, geographic distance, and elevation, using model selection based on AICc to identify the combination of factors that best explained variation in each phenotype. The best fit model explaining variation in temporal persistence among strains included significant effects of both elevation (β = 0.06 ± 0.01, t = 6.91, p < 0.001; Table S9) and neutral genetic variation (β = 1.19 ± 0.50, t = 2.38, p = 0.017), providing further evidence that this collective behavior has evolved in response to natural selection from elevation, while also confirming the expected strong role of neutral evolutionary processes. On the other hand, variation in aggregation magnitude was best explained by a model that only included neutral genetic variation (β = 2.12 ± 0.50, t = 4.34, p < 0.001;; Table S9). Taken together, these results isolate the expected effects of neutral evolutionary mechanisms on collective behavior trait variation to highlight additional support that aggregation temporal persistence has potentially evolved among *C. elegans* strains in response to natural selection from elevational differences experienced by different populations.

### Aggregation traits are heritable with a complex genetic basis

Next, we evaluated if aggregation behavior in *C. elegans* has the potential to respond to natural selection through genetic mediation. First, we tested how heritable the aggregation traits are by estimating the genetic variance from total behavioral variance using heritability calculations from CaeNDR. Using a linear mixed model to account for among-strain variation, we found that broad-sense heritability explains 61.82% (CI: [49.70%, 67.85%]) of the variation in mean aggregation magnitude and 36.02% (CI [19.50%, 45.31%]) of the variation in temporal persistence (Fig 4a top). Narrow-sense heritability was estimated by including the additive genetic information using the covariance of genotypes among strains, demonstrating that 54.96% (CI [42.69%, 62.28%]) of the variation in mean magnitude and 24.36% (CI [8.66%, 33.71%]) of the variation in temporal persistence are explained by the additive genetic effect (Fig 4b bottom). These heritability scores are greater than the average narrow-sense heritability of behavior traits (23.50%) (62), and suggest that natural variation in *C. elegans* aggregation has a genetic basis.

**Figure 4:**
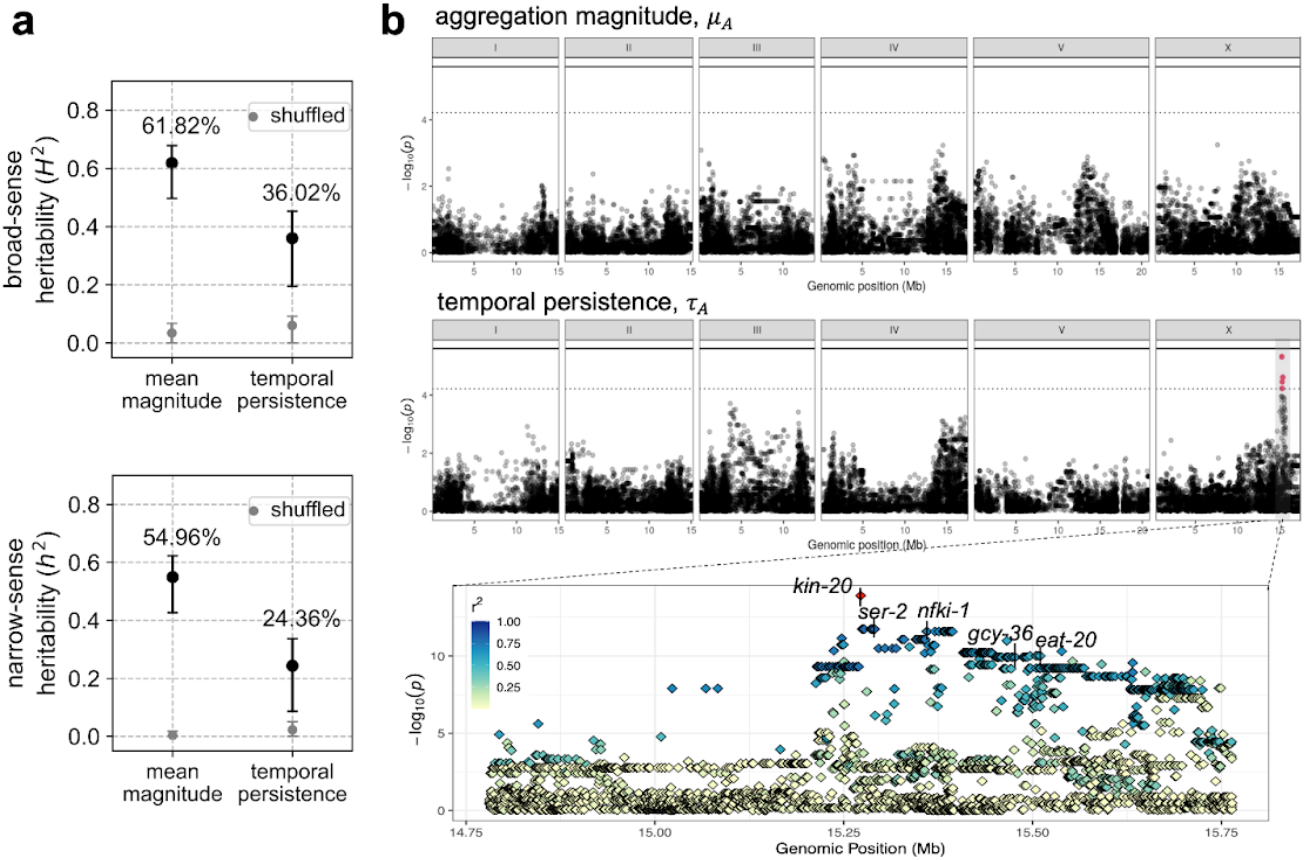
Genetic basis of the natural variation in aggregation behavior. a. Heritability of the aggregation traits. Broad-sense heritability assumes strains are independent, whereas narrow-sense considers the strains’ genetic covariances. Error bars denote 95% confidence interval of bootstrapped heritability. Grey points are control estimates for heritability generated by randomly shuffling across the strains. b. Top panel is the Manhattan plot for aggregation magnitude showing associations between genome-wide SNPs and the temporal persistence of aggregation. Middle panel is the Manhattan plot for temporal persistence. A QTL on chromosome X containing significant association is highlighted in red. The bottom panel zooms into the fine-mapping of that QTL. Colors depict *r*^2^ values, which denote the linkage of individual SNPs to the peak marker in red. Five candidate genes potentially associated with aggregation are labeled.

To further evaluate which genetic loci could underlie the observed behavioral variation, we performed GWAS where the associations between a phenotype and single nucleotide polymorphisms (SNPs) are assessed across the genome (63). GWAS mapping identified no genomic region significantly associated with aggregation magnitude (Fig 4b top, Fig S4). This lack of association could be simply due to a lack of statistical power arising from complex population structure found in selfing species (64), or to other reasons such as the polygenic nature of most complex traits precluding the identification of a single locus with a large effect, such as the case for human height (65, 66). GWAS for temporal persistence, on the other hand, reported a QTL on the right arm of chromosome X for this trait (Fig 4b top, Fig S6) across two mapping algorithms (see Methods). This QTL accounts for 39.58% of the additive genetic variance underlying this trait variation. Subsequent fine mapping within this QTL revealed a genomic region of around 430 kb in high linkage disequilibrium with the peak marker (Fig 4b bottom). There are eight non-coding regulatory RNAs and 25 protein-coding genes within this region, most of which function in basic cellular and developmental processes. Among them, we identified five candidate genes that could potentially modulate the temporal dynamics of *C. elegans* aggregation (Fig 4b bottom): *nfki-1* and *gcy-36* have been shown to affect aggregation behavior via oxygen response (67, 68) *kin-20* is a known regulator of rhythmic activity (69), and *ser-2* and *eat-20* modulate pharyngeal pumping and foraging behavior in *C. elegans* (70, 71) The linkage amongst these candidate genes suggests that the variation of aggregation temporal dynamics among the wild *C. elegans* strains may be genetically mediated via one or a combination of these genes. Interestingly, despite the strong causal effect of *npr-1* mutations on the aggregation phenotype in domesticated strains (44, 49–52), our GWAS results suggest that known natural variation in this gene has little effect on phenotypic variation in wild *C. elegans* populations.

## Discussion

We present *C. elegans* aggregation as a study system to address how collective behavior may evolve across natural habitats on a global scale. We screened a panel of 196 wild *C. elegans* strains sampled from around the world, and performed behavior analyses to reveal substantial natural variation in the mean magnitude and the temporal persistence of their aggregation behavior. We found that the elevation of the strain isolation sites predicts the observed behavioral variation, suggesting that elevation may impose selection pressure to drive this behavioral divergence. Moreover, both aggregation traits are highly heritable, and GWAS for temporal persistence revealed a QTL that may underlie phenotypic variation in this trait. Our results support that aggregation is a heritable collective behavior which was potentially under selection by local elevational conditions across diverse habitats.

A key development of our study is the experimental and analytical methods to effectively expose and measure natural variation in *C. elegans* aggregation behavior. Most previous studies of *C. elegans* aggregation treat it as a static and binary phenotype (44, 49–53). While semi-quantitative measurements were sufficient to show that wild *P. pacificus* populations have a clear binary divergence between solitary or aggregating phenotypes across an elevation threshold (53), the case with wild *C. elegans* is much less obvious and requires rigorous behavioral quantifications to reveal more subtle natural variation. Capturing such behavioral divergence allowed us to further examine the environmental and genetic associations to the behavior and make evolutionary inferences about *C elegans* collective behavior. Furthermore, a classic challenge for associating environmental and behavioural variation is that the environment can affect behaviour through both direct and indirect effects. For instance, temperature directly affects *C. elegans* speed and the wavelength of undulation (74), so that if aggregation were measured in the natural habitats at different temperatures, behavioral variation can be attributed to the direct temperature effect, confounding strain-level differences that may have evolved in different environments. Our controlled laboratory experiments with global populations thus dissociate the direct environmental effect on behavior and enable a clear interpretation of genetically mediated evolutionary effect.

Our results indicate that elevation is a strong environmental predictor of *C. elegans* aggregation behavior on a global scale. A similar pattern was found in the collective behavior of *P. pacificus* La Réunion island, where strains from high elevations show increased aggregation and vice versa. While the case with *P. pacificus* applies to a single clade, higher aggregation in our *C. elegans* dataset appears to have evolved multiple times across genetically distinct strains from higher elevations (Fig 3a). This suggests that elevational behavioral divergence has evolved several times in the nematode phylum (53), and points to a potential adaptive value of this behavior across different environments (73). The question remains of how elevation may drive the divergence of aggregation behavior. A leading hypothesis is differential physiological response in populations that have adapted to various ambient oxygen levels across elevations. A body of work in *C. elegans* suggests oxygen response as the main cause of aggregation in *npr-1* mutants, where the animals prefer low oxygen concentrations and cluster together to locally reduce O_2_ levels under the hyperoxic laboratory condition of 21% oxygen (44, 49–51). Here we showed that in wild populations, strains with higher aggregation in the lab indeed come from high elevation source habitats where atmospheric oxygen partial pressures are low. This is consistent with clustering for hyperoxic stress avoidance in these populations that have adapted to lower oxygen levels. Two genes identified from our candidate QTL, *nfki-1* and *gcy-36*, are known to modulate aggregation through oxygen sensory pathways (67, 68), also supporting that differential physiological responses may play a role in aggregation behavioral variation in wild *C. elegans* populations.

An alternative but not mutually exclusive hypothesis to the hyperoxia avoidance hypothesis is foraging. It has been suggested that lower oxygen concentrations may indicate actively proliferating and metabolizing bacterial sources for *C. elegans* to feed on (49, 50, 74). The presence of *ser-2* and *eat-20*, shown to regulate pharyngeal pumping and foraging, in our QTL further supports that foraging behavior in different resource environments could play a role in shaping aggregation phenotypes (70, 71) in wild *C. elegans*. Moreover, besides ambient oxygen level, elevation also correlates with other environmental conditions such as temperature fluctuations and microbial stability and composition (75–78), all of which may affect foraging behavior in natural *C. elegans* populations (79–81) Altered foraging behavior can plausibly influence aggregation: since animals in resource abundant environments are more likely to stay in their local search mode (81, 82), this could promote overall proximity between individuals and increase the probability of interaction and group formation. Therefore, the association between aggregation behavior and habitat elevation could potentially be explained by physiological hyperoxia avoidance, by a complex foraging-related response, by other abiotic and biotic factors with elevational variation that we have not yet considered (83), or by some combination of the above. Further work testing the physiology and the foraging hypotheses, such as by assessing the potential costs of being in hyperoxic environments and by examining aggregation dynamics under ‘native’ oxygen concentration in various food conditions, would help to pinpoint what is under selection and what is the potential adaptive value of this behavioral variation.

Additionally, the speculated role of hyperoxia avoidance and foraging behavior on emergent aggregation implies that variation in collective phenotypes may derive from the evolution of individual-level behavior. However, interactions between individuals often play an important role in shaping collective behavior (84) and could also be subjected to evolution. Future studies scrutinizing the individuals’ oxygen gradient response and foraging states as well as their interactions inside the aggregating group, in a comparative context across different wild populations, could help disentangle individual level versus inter-individual level mechanisms in the evolution of *C. elegans* collective behavior.

In summary, our study reveals natural variation in a collective behavior across global populations of *C. elegans* and identifies its environmental and genetic correlates, suggesting natural selection has shaped the evolution of this collective behavior. With this new evidence of collective behavior evolution, a collection of geo-referenced wild strains showing quantitative genetic and behavioral variation, and empirical possibilities for vigorous downstream validation, aggregation behavior in *C elegans* serves as a powerful system for future studies to reveal the evolutionary dynamics of collective behavior.

## Methods

### *C. elegans* strains

The *npr-1*(*ad609*) knock-out mutant strain was obtained from the *Caenorhabditis* Genetics Center (CGC). The N2 laboratory reference strain and the panel of 196 wild strains of *C. elegans* obtained from CaeNDR (54), with the list of wild strains in Table S7. All animals were regularly cultured and maintained on nematode growth media (NGM) plates using standard protocol (85), and fed *E. coli* OP50 as worm food. *E. coli* OP50 was obtained from the CGC and was cultured in Luria Broth using standard liquid culture protocol (86)

### Behavioral assay and video acquisition

Animals for the behavioral assay were prepared as synchronized Day-1 adults using standard bleaching protocol (87). 35 mm plates pre-filled with low-peptone NGM (88) were used for imaging, where the center of each plate was seeded with 75 µL of diluted OP50.A master batch of imaging OP50 was made for the entire dataset collection by diluting an overnight liquid culture 1:10 in M9 to a final concentration of OD_600_ = 0.384; aliquots from this were stored at 4 ºC until they were used to freshly seed the imaging plates on the day of the experiment. The entire food patch was within the field of view of each camera. Synchronized worms were collected from culture plates and washed twice in M9, and 40 individuals were transferred to the agar surface off food using a glass pipette. After M9 has dried, gentle vortexing was applied at 600 rpm (Vortex Genie 2 shaker, Scientific Industries) for 10 seconds to randomize initial positions and synchronize the aggregation state across replicates and strains. Imaging was performed immediately afterwards for 45 minutes at 25 Hz, with a custom-built six-camera array (Dalsa Genie Camera, G2-GM10-T2041) under red illumination (630 nm LED illumination, CCS Inc), maintaining 20 ºC throughout the experiment. Six independent experiments were run simultaneously and strain identity, camera position, and recording session assignments were randomized across experiments. Gecko software (v2.0.3.1) drove simultaneous data acquisition from the six cameras. At least five independent experimental replicates were performed for each strain, with N2, *npr-1*, CB4856 strains exceeding the number due to extra recordings taken as internal controls per day for every session. Experiments were inspected during and after data collection; experiments with puncture or debris in the agar were excluded from the dataset.

### Video processing and cluster size estimation

For post-acquisition video analysis, a custom algorithm was developed to estimate multi-worm cluster sizes from processed video recordings. Following background subtraction, images were binarized to detect discrete objects (blobs). Videos were acquired at 25 frames per second, with every third frame sampled for analysis (effective rate: 8.33 Hz). The approach operates on a graph-based representation where detected objects (blobs) in each frame serve as nodes, connected across consecutive frames based on minimal inter-component pixel distances. Single worms were identified through an iterative classification process that integrates morphological features (object area relative to dynamically computed thresholds) and trajectory spatial extent (the area of the polygon encompassing all positions along each object’s trajectory). This approach exploits the greater mobility of single worms, which exhibit larger trajectory areas compared to multi-worm clusters. Groups identified as singles based on these criteria—specifically those where a majority of trajectory frames fell below the area threshold and exhibited low in-degree and out-degree in the temporal graph—were assigned a size of one worm. For multi-worm clusters, size estimation proceeded through an iterative propagation algorithm operating on the temporal graph structure. At each iteration, the method identifies local subgraphs where cluster sizes can be uniquely determined from conservation constraints: when clusters split or merge between frames, and all but one resulting group size is known, the unknown size is computed from the difference in total worm counts. This process continues until convergence or a maximum iteration limit is reached. For clusters where size remains ambiguous after convergence, the mean of the minimum and maximum possible sizes was assigned. It should be noted that for large, frequently splitting and merging clusters, this approach may introduce upward bias in size estimates. Videos exhibiting tracking irregularities that produced non-continuous time series were excluded from downstream analyses.

### Aggregation trait measurements

To estimate the magnitude of aggregation at each time *t*, the inverse of the spatial entropy of the distribution of worms in each frame was computed. Cluster identities were treated as discrete bins whose probabilities were given by the estimated number of worms in each cluster divided by the total number of worms (6):

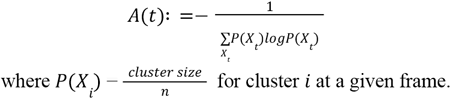

From the time series *A*(*t*) *t* ϵ {δ*t*,*…, T*δ*t*} obtained from each trial, where δ*t* is the inverse frame rate, we estimated mean magnitude as the average of the time series,

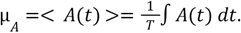

To capture the temporal dynamics of the time series *A*(*t*), the normalized autocorrelation function was measured,

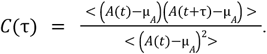

The complex nature of the correlation functions, which deviate from simple exponential decay (likely due to finite-size effects), challenges the inference of decay times for each trial (89, 90). To nonetheless capture the strength of temporal correlations, we define an overall measure of temporal persistence of the aggregation dynamics, τ_*A*_, as the maximum of the cumulative sum of the correlation function,

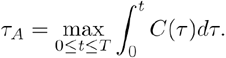

This measure is analogous to the integral timescale used in the studies of fluids and turbulence, which assesses the overall temporal memory of the dynamics (91). To account for the imbalance in sample size between CB4856 and other wild strains for the environmental and genetic analyses, bootstrapping of *μ*_*A*_ and τ_*A*_ values across replicates was performed to achieve an equal final number of five samples per strain.

### Environmental data

Using the GPS coordinates of the isolation sites of each wild strain, four environmental variables for the PGLS analysis were obtained from two public databases. Elevation data was obtained from the OpenElevation API (https://open-elevation.com). Climate variable data including near-surface temperature, humidity, and upper column soil moisture were obtained from the CMIP6 global climate model (92, 93) of the Copernicus Climate Change Service (C3S) Climate Data Store (DOI: 10.24381/cds.c866074c); specific model details are in Table S1 (see Data availability). Monthly data from years 2000-2014 was obtained to cover the relevant isolation period for all of our wild strains except CB4856, which was collected in 1972 before the CMIP6 climate model data began. Given the periodicity of the climate variables and the simplicity of the measurements (mean and standard deviation), we used the 2000-2014 data to approximate 1972 for this one strain. Each climate variable was interpolated into spatial resolution of 1.4° latitude and longitude spanning the entire globe. Isolation sites of the wild strains were matched to geographical coordinates with less than 0.7° error in both latitude and longitude. Out of the 196 wild strains, five strains (CB4852, ECA252, ECA259, LSJ1, PB303) missing the GPS coordinates and one (JU2001) missing the climate information from the CMIP6 database were excluded from the analysis.

### Generalised Least Squares estimation of aggregation traits

In comparative evolutionary biology assessing phenotypes between populations, a generalised linear model (GLM) is used to account for phylogenetic relationship to avoid the assumption of data independence and overestimation of statistical significance (94) (95, 96). Therefore, a linear mixed model with phylogenetic inter-strain covariance as the random effect was used to estimate the true effects of predictors for our wild population behavioral phenotypes. The linear mixed model is:

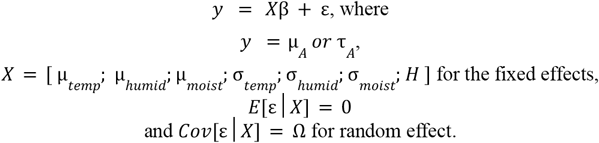

Parameter β of intercept and slopes is estimated based on the derivation of generalised least squares (GLS) regression:

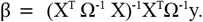

The resulting phenotype accounted for the inter-strain covariance is computed as:

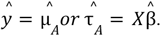

The nlme (97) and ape (98) packages of R were used to compute the GLS. Correlation structure of the residual Ω was specified as the Brownian motion process of evolution, by the function corBrownian of the ape package (94)

### Partitioning the effects of natural selection and neutral evolution on aggregation traits

Maximum likelihood population effects (MLPE) models were used to test for the relative roles of neutral evolution (genetic drift, gene flow) and natural selection in shaping the observed variation in aggregation behaviors. MLPEs are mixed effect models used to evaluate the relationship between two or more pairwise distance matrices for all combinations of populations, including a population effect to account for the nonindependence of pairwise distance comparisons (99). If aggregation behaviors are the result of natural selection from local environmental conditions, an association between phenotypic variation and environmental variation is expected to be observed after controlling for the effects of genetics and geography, reflecting that populations are more phenotypically similar if they come from similar habitats, regardless of their genetic or geographic proximity (100). If aggregation behaviors have been shaped by neutral processes, phenotypic distance should show a significant association with geographic and/or genetic distances, indicating that populations that are closer together or genetically more similar are more phenotypically similar. We consider this scenario simply as “neutral evolution,” because phenotypic similarity could arise from gene flow between geographically proximate populations as well as neutral evolution resulting from genetic drift between populations geographically isolated from each other (100).

Pairwise phenotypic distances between populations were estimated separately for both aggregation magnitude and temporal persistence. Each trait was standardized via z-score transformation with the scale function in base R before calculating the Euclidean distances between every pairwise population combination using the population mean trait value with the dist function from the stats package (101) Pairwise Euclidean distances between standardized elevation values for each collection site were calculated in the same manner. Pairwise geographic distances between each collection site were calculated as the geodesic distance in kilometers between coordinates with the geodist package (102) To facilitate comparisons with the other distance matrices, geographic distances were scaled by dividing each pairwise value by the maximum distance in our matrix. Genome-wide genotype data from the 20250625 CaeNDR (54) release was used to estimate genetic distances between populations. Specifically, the hard-filtered variants with imputed missing genotypes (WI.20250625.impute.isotype.vcf.gz) were downloaded for the 190 isolines with complete phenotype and environmental data. To obtain a set of putatively neutral SNPs across the genome, bcftools (v.1.23; (103)) was used to remove all SNPs found within hyper-divergent regions in the *C. elegans* genome (39), as these regions are characterized by extreme sequence divergence between strains and are enriched for genes hypothesized to play a role in local adaptation (104). This SNP set was further pruned by removing sites in linkage disequilibrium using PLINK (v1.9; (105)), using the --indep-pairwise command to identify and prune variants with an ^*2*^ value greater than 0.2 in 50 kbp windows, sliding forward 5 variants after each pruning step and repeating. Filtering and pruning resulted in a final VCF file containing 255,853 biallelic markers, which was used to estimate pairwise genetic distances between all population pairs calculated as the proportion of nucleotide sites at which the two genomes differ (*p*-distance) using VCF2Dis (v1.55; (106)). To assess the role of selection and neutral evolution in shaping aggregation behavior, MLPE models were fit using corMLPE (https://github.com/nspope/corMLPE) and the *nlme* package in R (97, 107), and the best fit model was selected based on the Akaike Information Criterion corrected for finite sample size (AICc) using the *MuMIn* R package (108). First, the genetic distance matrix was confirmed to be representative of neutral genetic variation among populations by fitting models using the pairwise genetic distance estimates as a response variable with either the elevation distance matrix, geographic distance matrix, or both matrices as fixed effects and population pairs modeled as random effects. To test the relationship between behavioral distances, natural selection, and neutral evolution, models were separately fit with either aggregation spatial magnitude or temporal persistence as a response variable and genetic distance as an explanatory variable, with population pairs included as random effects. Additional explanatory variables were added to this model to determine their relative effects, fitting models with either elevational distance, geographic distance, or both together.

### Heritability estimation

Broad-sense heritability ((^2^) and narrow-sense heritability (*h*^2^) were estimated from a subset of 12 genetically diverse strains (N2, CB4856, CX11314, DL238, ED3017, EG4725, JT11398, JU258, JU775, LKC34, MY16, MY23) (54) to assess the respective strain-group or additive genetic effect on the behavioral traits. The linear mixed model is,

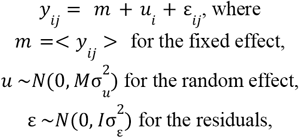

And *y*_*i*_ = [*y*_*i*1_ … *y*_*ij*_], ε = [ε _*i*1_… ε _*ij*_], for *i* = strain number (n), and *j* = replicate number. The R sommer package was used to estimate 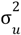 and 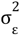 by maximizing the log likelihood function computed based on the Restricted Maximum Likelihood (REML) of the *Y* distribution (109). Heritability, 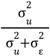 captures the ratio of strain or additive genetic variance over total phenotypic variance. The model is inferred with M=I (the identity matrix) for (^2^ and M=A for *h*^2^. The additive genetic relations matrix was constructed using the Van Raden (110) method, 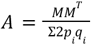, where the covariance of the genotypes among strains (*MM*^*T*^) at different SNPs is normalized by allele frequency (*p*_*i*_ = frequency of allele 1 at locus *i* and *q*_*i*_ = frequency of allele 2 at locus *i*) (54, 111, 112)

The code for estimating the heritability was adapted from the CaeNDR heritability calculation (https://github.com/AndersenLab/calc_heritability.git). Main dependencies include the sommer package (113) of R and bcftools (114) to generate the genotype matrix from the genome sequence of the strains (see Code Availability).

### Genome wide association analysis

To find genetic correlates of the aggregation traits, GWAS analyses were performed to assess the associations between variation in phenotypic traits and naturally occurring genetic variation among the wild strains (115, 116). The NemaScan pipeline implemented by CaeNDR was used with default parameters; the pipeline accounts for the specific genetic architecture of the species that could bias the mapping performance, such as the recent genome-wide selective sweep in *C. elegans* and population stratification among self-fertilizing populations (64). The pipeline scans through the chromosomes and prunes genetic variants with high linkage disequilibrium in 50 Kb windows, then computes the associations between retained genetic variants and phenotypic values (64). If a variant with high associations indicative of a Quantitative Trait Locus (QTL) is returned, subsequent fine mapping specifies the associations of all single nucleotide polymorphism markers within the QTL region (64) to the trait. Fine-mapped variant markers had 250-600 bp intervals on average. The two mapping algorithms of NemaScan consists of two kinship matrix formulations: a leave-one-chromosome-out (LOCO) and the ‘INBRED’ approach that specifically corrects for inbred organisms (64) such as the self-fertilizing *C. elegans*. Both mappings attempt to correct for different types of population structure found in *C. elegans* (64). For the significance threshold, two metrics were reported: a Bonferroni score correcting for the total number of all tested variants, and an ‘EIGEN’ score that performs Bonferroni correction with the number of tests determined by the eigendecomposition of the genetic variants (64) Input data such as the hardfiltered SNPs, haplotypes, and the hyper-divergent regions of the strains were downloaded from the 20250625 CaeNDR release (https://www.elegansvariation.org/data/release/20250625). Four strains from the global panel (ECA252, JU1580, LSJ1, QX1233) were missing from this data.

To scrutinize the effect of population structure on GWAS significance, outlier strains were additionally pruned following detection by smartPCA implemented in the *smartsnp* package of R (117), which allows centering and z-score standardization on genotype matrix to correct for genetic drift and population structure (117, 118). GWAS was rerun after excluding the 10 outlier strains above the 97.45% Mahalanobis threshold, and the results were confirmed to maintain the same overall pattern (Fig S5, S7).

### QTL variance estimation

The variance explained by the QTL was estimated as the ratio of QTL variance over total phenotypic variance,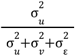. (119, 120). The linear mixed model that estimates QTL variance separately from the rest of the additive genetic effect was used:

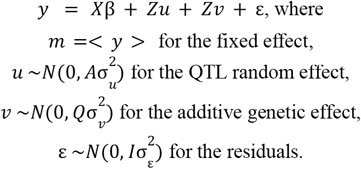

Genetic relations matrix for the QTL region was constructed using the same Van Raden method 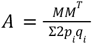 (110), but only among the SNPs within the QTL. Similarly, the genetic relations matrix *Q* was computed as the covariance of the genotypes among strains excluding the SNPs within the QTL (111, 112, 120)

## Data Availability

Datasets used in this study will be available at Zenodo upon publication (https://zenodo.org/uploads/14937989), which includes original tracking data from the experiments in this study, climate data obtained from Climate Data Store, information on the natural habitats of the wild *C. elegans* strains, and phylogenetic information from CaeNDR.

## Code Availability

Code for analyses and generating figures are deposited on Github (https://github.com/SerenaDingLab/Kang_et_al_AggEvol25).

## Acknowledgements and Funding Information

We are grateful for Erik Andersen for his helpful insights and comments on wild *C. elegans* GWAS analysis. We thank *Caenorhabditis* Natural Diversity Resources for the wild *C. elegans* strains and the genetic analysis tools. Additional strains were provided by the CGC, which is funded by NIH Office of Research Infrastructure Programs (P40 OD010440). We acknowledge the World Climate Research Programme and its Working Group on Coupled Modelling who coordinated and promoted CMIP6. We also thank multiple funding agencies who support CMIP6 and ESGF, and the climate modeling groups and the Earth System Grid Federation (ESGF) for producing and archiving the data. This research was supported by the Max Planck Institute of Animal Behavior, the International Max Planck Research School for Quantitative Behavior, Ecology & Evolution, the Human Frontier Science Program (RGP0001/2019), the Medical Research Council through grant MC-A658-5TY30, and the Deutsche Forschungsgemeinschaft (462886202). We also thank Claire Wyart for the support, and acknowledge funding to A.C.C. from the Agence Nationale pour la Recherche LOCOCONNECT and RSCMAP awarded to C.W. An open access license has been selected for this work.

## Author Contributions

Y. J. K., A.E.X.B. and S. S. D. conceptualized the project; S. S. D. and P. S. performed the experiments; Y. J. K., A. C. C. and A.P. analyzed the behavior data; Y. J. K. and R. G. analyzed the genetic data; Y. J. K. prepared the initial manuscript; Y. J. K., A. C. C., R. G., A. P., A.E.X.B. and S. S. D. reviewed and edited the manuscript; S. S. D. and A.E.X.B. supervised the project and acquired funding.

