## Supplementary Information for "Evidence for natural selection shaping the evolution of collective behavior among global *Caenorhabditis elegans* populations"

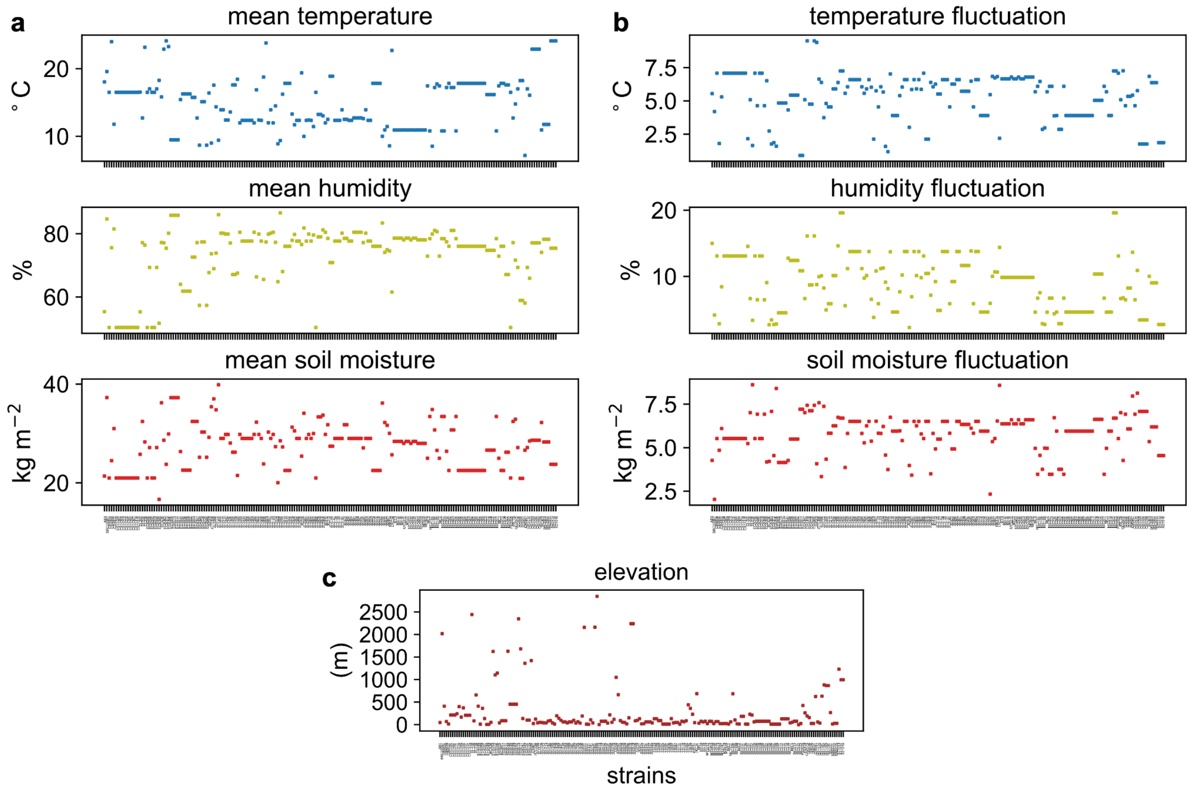

**[Fig S1] Environmental information of the strain isolation sites**. a. Near-surface temperature, near-surface relative humidity and upper column soil moisture of the isolation sites of the 190 wild strains, time-averaged over the period 2000-2014. b. Standard deviation of the temperature, relative humidity and soil moisture of the isolation sites of the wild strains, over the period 2000-2014 . c. Elevation of the isolation sites of the wild strains.

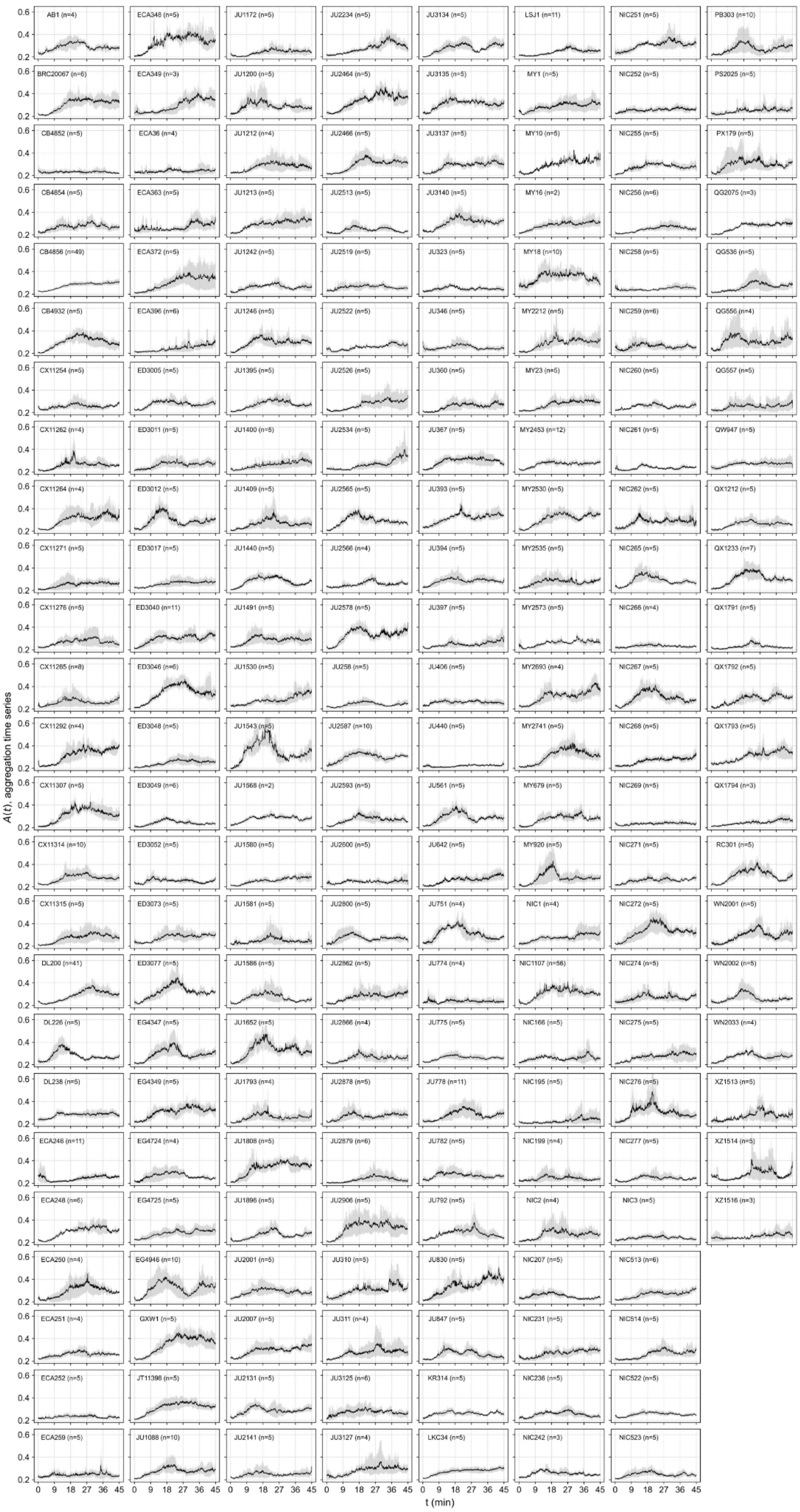

**[Fig S2] Natural variation in aggregation time series**. Aggregation time series of the 196 wild strains over 45 minutes. Line depicts a strain's aggregation magnitude averaged across experimental replicates, and shading is the 95% bootstrapped confidence interval to capture experimental variation.

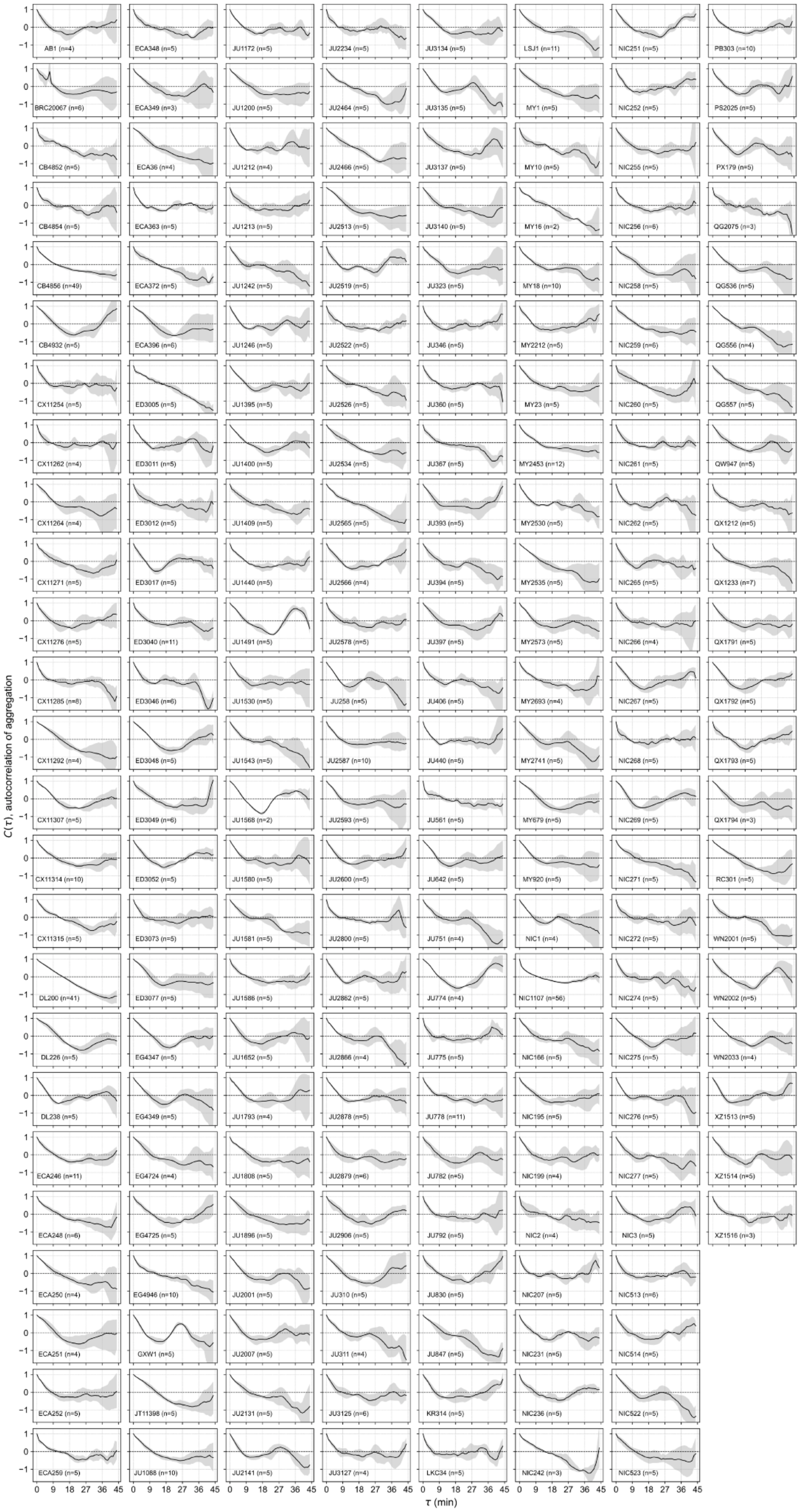

**[Fig S3] Natural variation in aggregation dynamics**. Autocorrelation function of the aggregation time series of the 196 wild strains as the measure of aggregation dynamics. Line depicts a strain's autocorrelation function averaged across experimental replicates, and shading is the 95% bootstrapped confidence interval to capture experimental variation.

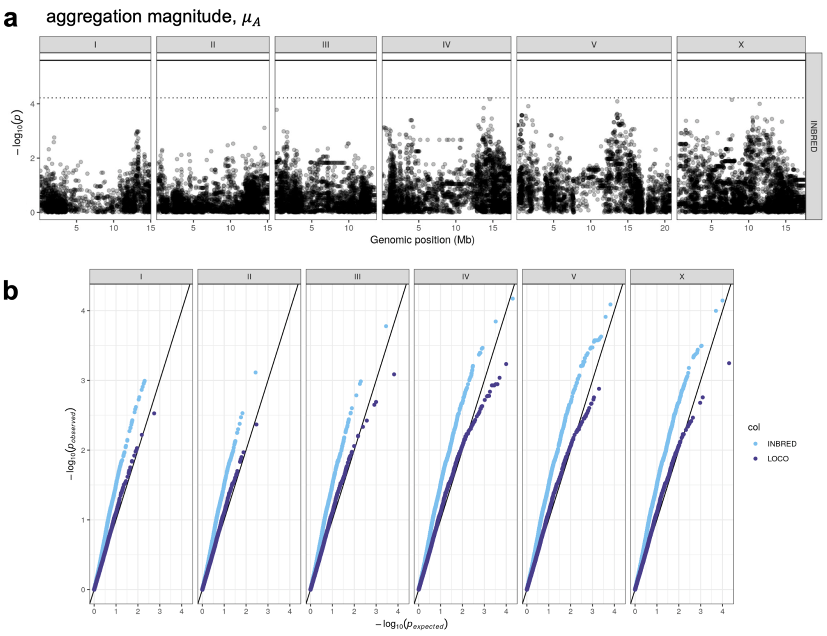

**[Fig S4] GWAS result of the aggregation magnitude.** a. Manhattan plot of aggregation magnitude showing genetic variants associated with natural variation in the phenotype (INBRED mapping). There is no genetic variant that surpassed the significant threshold. Dotted line refers to the EIGEN threshold while bold line refers to the Bonferroni score. b. Q-Q plot depicting the Genomic inflation factor of the INBRED and LOCO mapping. Solid line represents when the p-value obtained from the analysis matches the theoretical $\chi^2$ p-values. Genomic inflation factor was 1.8031051 for the INBRED mapping and 1.1304331 for the LOCO mapping.

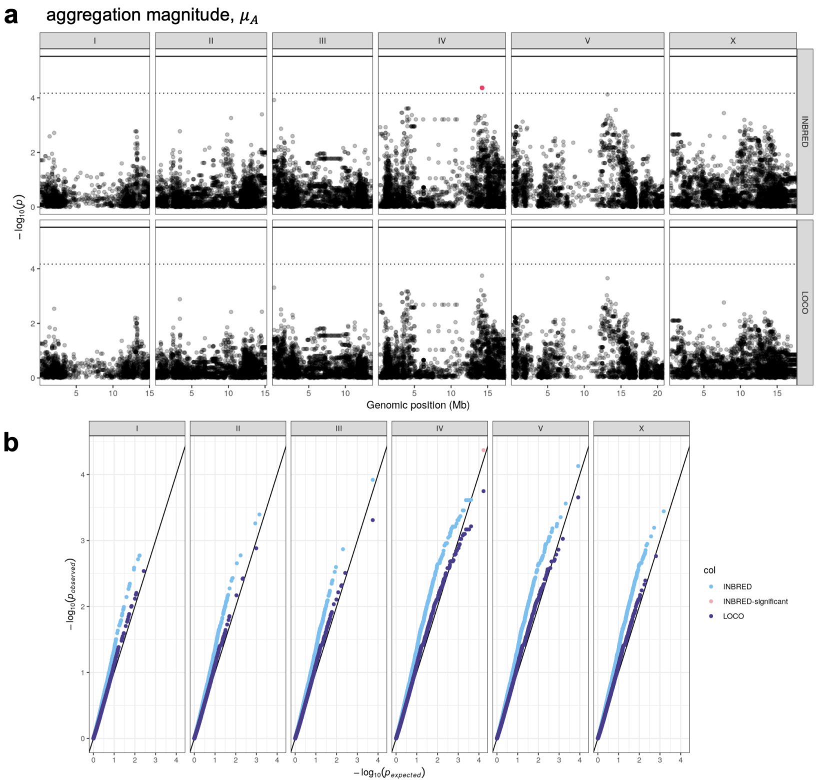

**[Fig S5] GWAS result of the aggregation magnitude after pruning for population stratification using smartPCA.** a. Manhattan plot of aggregation magnitude after pruning the outlier strains from smartPCA. Top panel refers to the result of the EIGEN algorithm and bottom panel the LOCO algorithm. Red dot refers to the peak marker of the genetic variant that surpassed the significant threshold of EIGEN score. b. Q-Q plot depicting the Genomic inflation factor of 1.4321477 for the INBRED mapping and 1.0482342 for the LOCO mapping.

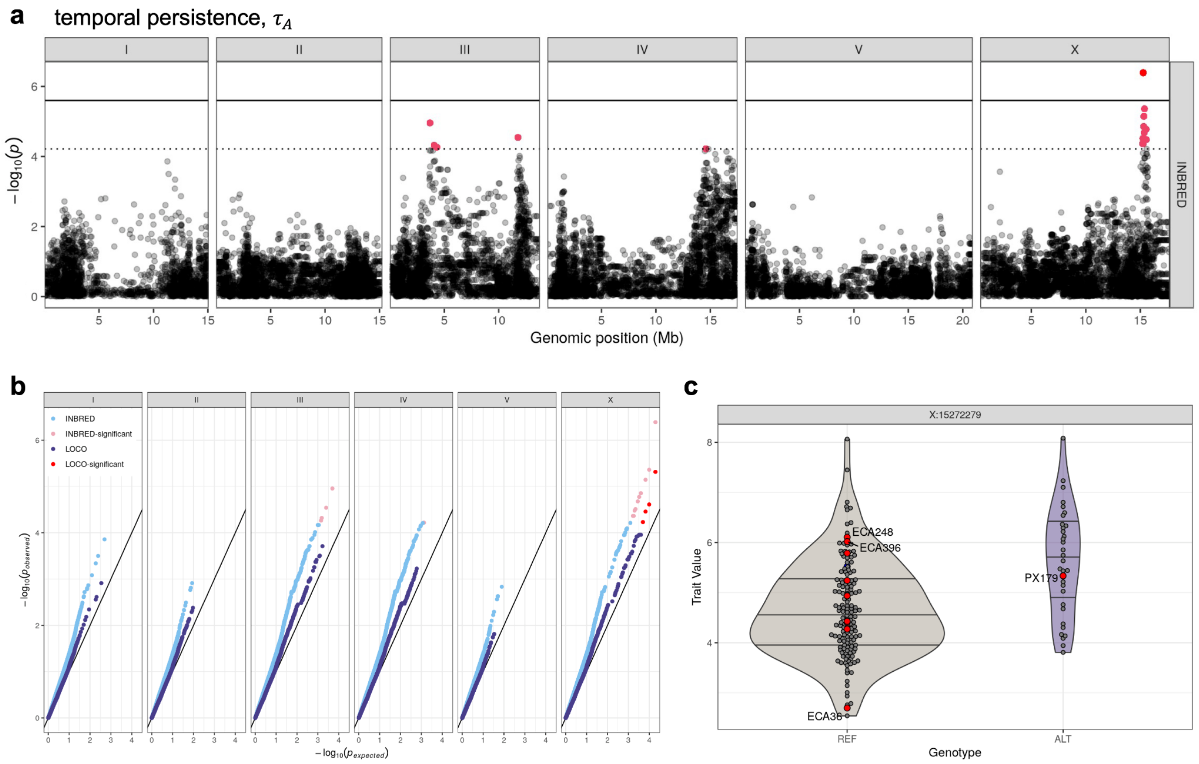

**[Fig S6] GWAS result of the aggregation temporal persistence.** a. Manhattan plot of aggregation temporal persistence showing the genetic variants associated with natural variation in the phenotype (INBRED mapping). Red dots refer to the peak markers of the genetic variants that surpassed the significant threshold of EIGEN score. Dotted line refers to the EIGEN threshold while bold line refers to the Bonferroni score. b. Q-Q plot depicting the Genomic inflation factors of the INBRED and LOCO mapping. Solid line represents when the p-value obtained from the analysis matches the theoretical $\chi^2$ p-values. Genomic inflation factor was 1.6994635 for the INBRED mapping and 1.1259024 for the LOCO mapping. c. Temporal persistence trait split into two groups based on the genotypes at the peak marker on chromosome X. Some strains with high genetic divergence as defined by CaeNDR are annotated as red points. No outlier strains driving the divergence are observed, suggesting that the associations found are not driven by a few outlier strains.

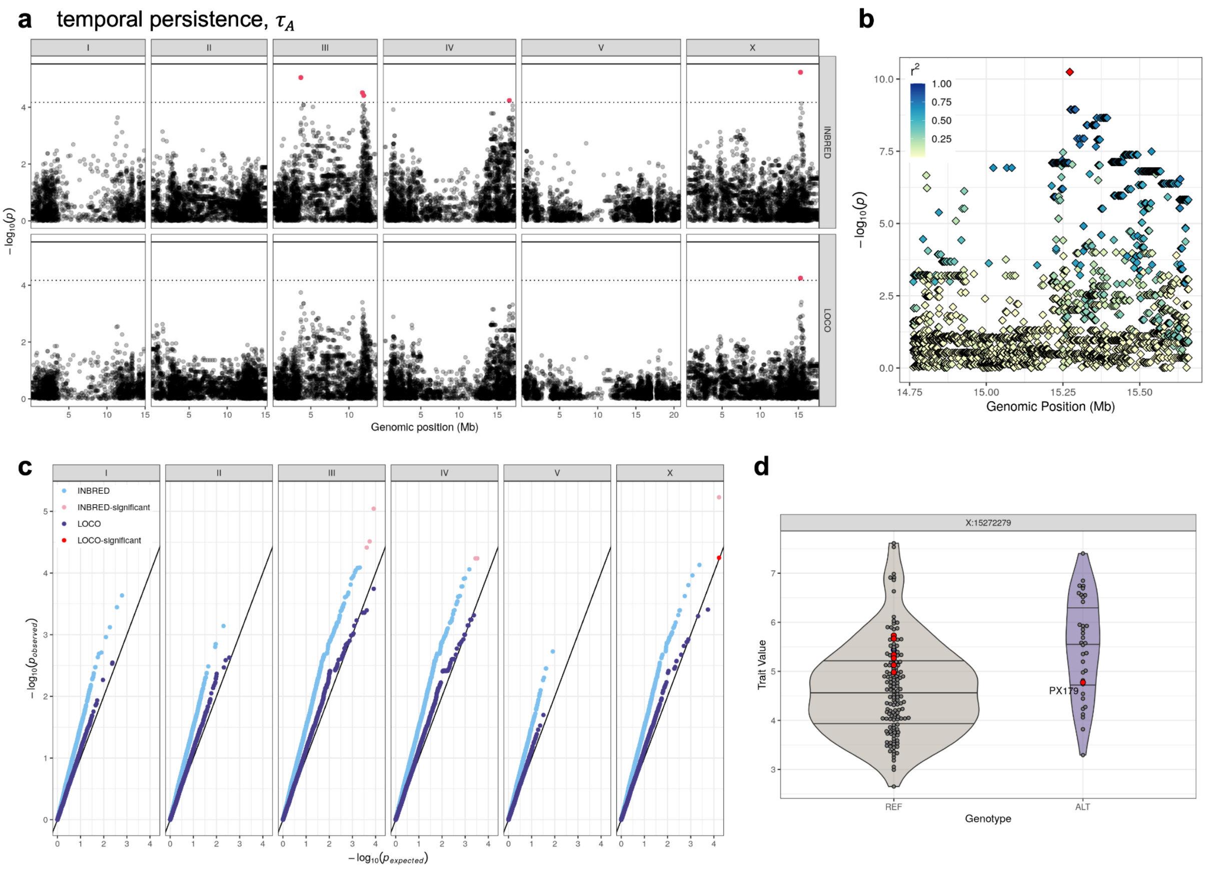

**[Fig S7] GWAS result of the aggregation temporal persistence after pruning for population stratification using smartPCA.** a. Manhattan plot of aggregation temporal persistence showing the genetic variants associated with natural variation in the phenotype. Top panel refers to the result of the EIGEN algorithm and the bottom panel the LOCO algorithm. Red dots refer to the peak markers of the genetic variants that surpassed the significant threshold of EIGEN score. Dotted line refers to the EIGEN threshold while bold line refers to the Bonferroni score. One QTL on chromosome X reports significant association in both the INBRED and LOCO mapping. c. Fine mapping result showing the SNP resolution mapping within the QTL at chromosome X. $r^2$ denotes the level of linkage to the red peak marker. c. Q-Q plot depicting the Genomic inflation factors of the INBRED and LOCO mapping. Solid line represents when the p-value obtained from the analysis matches the theoretical $\chi^2$ p-values. Genomic inflation factor was 1.9977412 for the INBRED mapping and 1.1747531 for the LOCO mapping. d. Temporal persistence trait split into two groups based on the genotypes at the peak marker on chromosome X. Some strains with high genetic divergence as defined by CaeNDR are annotated as red points. No outlier strains driving the divergence are observed, suggesting that the associations found are not driven by a few outlier strains.

**[Table S1] Source information of the CMIP6 climate model**

| source ID | institution ID | citation |
|---|---|---|
| CNRM-ESM2-1 | CNRM-CERFACS | Seferian, Roland (2018). CNRM-CERFACS CNRM-ESM2-1 model output prepared for CMIP6 CMIP historical. Version 20241215. Earth System Grid Federation. https://doi.org/10.22033/ESGF/CMIP6.4068 |

**[Table S2] Phylogenetic GLS aggregation magnitude model result**

| | Value | Std.Error | t-value | p-value |
|---|---|---|---|---|
| (Intercept) | 0.0056946 | 0.08070428 | 0.0705615 | 0.9438 |
| $H_{elevation}$ | 0.0606059 | 0.02573341 | 2.3551435 | 0.0196 |
| $\mu_{temp}$ | -0.0956418 | 0.13703497 | -0.697937 | 0.4861 |
| $\mu_{humid}$ | -0.0139916 | 0.08374576 | -0.167072 | 0.8675 |
| $\mu_{moist}$ | 0.0464189 | 0.13734817 | 0.3379652 | 0.7358 |
| $\sigma_{temp}$ | 0.346845 | 0.18223536 | 1.9032805 | 0.0586 |
| $\sigma_{humid}$ | -0.1948629 | 0.19062997 | -1.0222048 | 0.308 |
| $\sigma_{moist}$ | 0.0643088 | 0.0558555 | 1.1513428 | 0.2511 |

**[Table S3] Model selection on phylogenetic GLS aggregation magnitude**

| model | AIC |
|---|---|
| $\mu_A \sim \sigma_{temp} + H_{elevation}$ | 412.9324 |
| $\mu_A \sim \mu_{temp} + \sigma_{temp} + H_{elevation}$ | 413.2035 |
| $\mu_A \sim \mu_{moist} + \sigma_{temp} + H_{elevation}$ | 413.8245 |
| $\mu_A \sim \mu_{humid} + \sigma_{temp} + H_{elevation}$ | 414.1748 |
| $\mu_A \sim \sigma_{temp} + \sigma_{moist} + H_{elevation}$ | 414.4732 |
| $\mu_A \sim \mu_{temp} + \sigma_{temp} + \sigma_{moist} + H_{elevation}$ | 414.6676 |

**[Table S4] Phylogenetic GLS temporal persistence model result**

| | Value | Std.Error | t-value | p-value |
|---|---|---|---|---|
| (Intercept) | 0.0291098 | 0.0768041 | 0.379014 | 0.7051 |
| $H_{elevation}$ | 0.09216283 | 0.02448979 | 3.763316 | 0.0002 |
| $\mu_{temp}$ | 0.06999761 | 0.13041251 | 0.53674 | 0.5921 |
| $\mu_{humid}$ | -0.0350763 | 0.07969859 | -0.440112 | 0.6604 |
| $\mu_{moist}$ | -0.0172041 | 0.13071057 | -0.13162 | 0.8954 |
| $\sigma_{temp}$ | 0.29121683 | 0.17342851 | 1.679175 | 0.0948 |
| $\sigma_{humid}$ | -0.2785657 | 0.18141743 | -1.535496 | 0.1264 |
| $\sigma_{moist}$ | 0.06991003 | 0.05315619 | 1.315181 | 0.1901 |

**[Table S5] Model selection on phylogenetic GLS temporal persistence**

| model | AIC |
|---|---|
| $\tau_A \sim \mu_{humid} + H_{elevation}$ | 393.0655 |
| $\tau_A \sim H_{elevation}$ | 393.2248 |
| $\tau_A \sim \sigma_{temp} + H_{elevation}$ | 393.6564 |
| $\tau_A \sim \mu_{moist} + H_{elevation}$ | 394.269 |
| $\tau_A \sim \mu_{temp} + \sigma_{temp} + H_{elevation}$ | 394.4203 |
| $\tau_A \sim \mu_{humid} + \sigma_{temp} + H_{elevation}$ | 394.5468 |

**[Table S6] Covariance among GLS model predictors**

| | $\mu_{temp}$ | $\mu_{humid}$ | $\mu_{moist}$ | $\sigma_{temp}$ | $\sigma_{humid}$ | $\sigma_{moist}$ | $H_{elevation}$ |
|---|---|---|---|---|---|---|---|
| $\mu_{temp}$ | 1 | -0.2793096 | -0.5081034 | -0.5486109 | -0.3747677 | 0.03813078 | 0.143063 |
| $\mu_{humid}$ | -0.2793096 | 1 | 0.65387556 | -0.306686 | -0.450045 | -0.0463122 | -0.0963534 |
| $\mu_{moist}$ | -0.5081034 | 0.65387555 | 1 | -0.004855 | -0.104441 | -0.1541691 | 0.04301633 |
| $\sigma_{temp}$ | -0.5486109 | -0.306686 | -0.004855 | 1 | 0.65530001 | 0.00746113 | -0.4870412 |

| | | | | | | | |
|---|---|---|---|---|---|---|---|
| $\sigma_{humid}$ | -0.3747677 | -0.450045 | -0.104441 | 0.65530001 | 1 | 0.31160359 | -0.0145277 |
| $\sigma_{moist}$ | 0.03813078 | -0.0463122 | -0.1541691 | 0.00746113 | 0.31160359 | 1 | -0.0688574 |
| $H_{elevation}$ | 0.143063 | -0.0963534 | 0.04301633 | -0.4870412 | -0.0145277 | -0.0688574 | 1 |

**[Table S7] Geographic coordinates and elevation of the isolation sites of wild *C. elegans* strains**

| strain | latitude | longitude | elevation(m) |
|---|---|---|---|
| AB1 | -34.93 | 138.59 | 48 |
| BRC 20067 | 24.072954 | 121.169736 | 2018 |
| CB4854 | 34.189 | -118.131 | 409 |
| CB4856 | 21.33 | -157.86 | 69 |
| CB4932 | 51.02 | -3.10000000000002 | 15 |
| CX11254 | 34.130138 | -118.114835 | 214 |
| CX11262 | 34.130138 | -118.114835 | 214 |
| CX11264 | 34.130138 | -118.114835 | 214 |
| CX11271 | 34.13712 | -118.12532 | 241 |
| CX11276 | 34.20111 | -118.21198 | 399 |
| CX11285 | 34.14331 | -118.05496 | 166 |
| CX11292 | 34.13531 | -118.30582 | 372 |
| CX11307 | 34.12946 | -118.10987 | 208 |
| CX11314 | 34.12946 | -118.10987 | 208 |
| CX11315 | 34.12946 | -118.10987 | 208 |
| DL200 | 9.03 | 38.74 | 2442 |
| DL226 | 44.5633 | -123.2821 | 83 |
| DL238 | 19.11 | -155.81 | 659 |
| ECA246 | 34.189 | -118.131 | 409 |
| ECA248 | 37.44 | -122.14 | 10 |
| ECA250 | 34.096 | -117.719 | 362 |
| ECA251 | 34.1 | -118.1 | 136 |
| ECA348 | 37.775334 | -122.253889 | 4 |
| ECA349 | 32.765758 | -117.236741 | 5 |
| ECA36 | -36.893333 | 174.745529 | 51 |
| ECA363 | 20.723611 | -156.304444 | 1623 |
| ECA372 | 22.121667 | -159.664444 | 1102 |
| ECA396 | 19.449831 | -155.235493 | 1144 |
| ED3005 | 55.94 | -3.36000000000001 | 39 |
| ED3011 | 55.92 | -3.19 | 85 |

| | | | |
|---|---|---|---|
| ED3012 | 55.92 | -3.19 | 85 |
| ED3017 | 55.92 | -3.19 | 85 |
| ED3040 | -26.166667 | 28.016667 | 1629 |
| ED3046 | -33.366667 | 19.316667 | 454 |
| ED3048 | -33.366667 | 19.316667 | 454 |
| ED3049 | -33.366667 | 19.316667 | 454 |
| ED3052 | -33.366667 | 19.316667 | 454 |
| ED3073 | -1.083333 | 36.65 | 2346 |
| ED3077 | -1.316667 | 36.8 | 1681 |
| EG4347 | 44.04789 | -123.07108 | 136 |
| EG4349 | 40.771467 | -111.87316 | 1360 |
| EG4724 | 41.6288 | -8.3476 | 99 |
| EG4725 | 41.6288 | -8.3476 | 99 |
| EG4946 | 40.72596 | -111.82184 | 1420 |
| GXW1 | 30.542889 | 114.419828 | 23 |
| JT11398 | 47.763944 | -122.275484 | 124 |
| JU1088 | 34.7613 | 138.0149 | 40 |
| JU1172 | -36.87 | -73.04 | 51 |
| JU1200 | 55.577 | -4.60000000000002 | 48 |
| JU1212 | 48.71 | -3.81 | 39 |
| JU1213 | 48.71 | -3.81 | 39 |
| JU1242 | 49.1269 | 1.9595 | 81 |
| JU1246 | 49.12618 | 1.96152 | 88 |
| JU1395 | 47.2199 | 0.04619 | 38 |
| JU1400 | 37.3845 | -5.988 | 14 |
| JU1409 | 37.468 | -5.637 | 196 |
| JU1440 | 41.41307 | 2.15231 | 142 |
| JU1491 | 46.63 | 1.06 | 96 |
| JU1530 | 48.7015 | 2.1725 | 62 |
| JU1543 | 48.7015 | 2.1725 | 62 |
| JU1568 | 48.8092 | 2.3862 | 40 |

| | | | |
|---|---|---|---|
| JU1580 | 48.7015 | 2.1725 | 62 |
| JU1581 | 48.7015 | 2.1725 | 62 |
| JU1586 | 46.63 | 1.06 | 96 |
| JU1652 | -34.86 | -56.19 | 35 |
| JU1793 | 49.12604 | 1.95114 | 66 |
| JU1808 | 48.81257 | 2.395372 | 34 |
| JU1896 | 37.999722 | 23.749673 | 191 |
| JU2001 | -21.1 | 55.5 | 2158 |
| JU2007 | 50.7608 | -1.32413000000003 | 9 |
| JU2131 | 48.0497 | -4.70499999999998 | 6 |
| JU2141 | 46.6293 | 1.0512 | 107 |
| JU2234 | 47.227 | -1.58300000000003 | 38 |
| JU2464 | -13.155 | -72.525 | 2162 |
| JU2466 | -13.257 | -72.266 | 2845 |
| JU2513 | 38.26016 | 22.0736 | 2 |
| JU2519 | 38.7175 | -9.14859999999999 | 77 |
| JU2522 | 38.7175 | -9.14859999999999 | 77 |
| JU2526 | 38.7175 | -9.14859999999999 | 77 |
| JU2534 | 49.357921 | 0.097087 | 34 |
| JU2565 | 44.958394 | 1.066213 | 218 |
| JU2566 | 48.83886 | 2.44599 | 52 |
| JU2578 | 48.87613 | 2.4315 | 118 |
| JU258 | 32.73 | -16.89 | 1049 |
| JU2587 | 45.026713 | 3.872016 | 664 |
| JU2593 | 48.822556 | 2.222334 | 88 |
| JU2600 | 49.1242 | 1.9496 | 54 |
| JU2800 | 48.79 | 2.331 | 49 |
| JU2862 | 52.1944 | 0.1264 | 17 |
| JU2866 | 34.0751 | -118.4409 | 152 |
| JU2878 | 19.29732 | -99.098769 | 2240 |
| JU2879 | 19.29732 | -99.098769 | 2240 |

| | | | |
|---|---|---|---|
| JU2906 | 43.8023 | 11.2806 | 84 |
| JU310 | 46.63 | 1.06 | 96 |
| JU311 | 44.42 | 4.4 | 132 |
| JU3125 | 38.1127 | 13.3722 | 17 |
| JU3127 | 38.11196 | 13.37274 | 20 |
| JU3134 | 48.7015 | 2.1725 | 62 |
| JU3135 | 48.7015 | 2.1725 | 62 |
| JU3137 | 48.7015 | 2.1725 | 62 |
| JU3140 | 48.71 | -3.81 | 39 |
| JU323 | 44.42 | 4.4 | 132 |
| JU346 | 44.42 | 4.4 | 132 |
| JU360 | 48.98 | 2.23 | 90 |
| JU367 | 48.98 | 2.23 | 90 |
| JU393 | 49.28 | -0.319999999999993 | 13 |
| JU394 | 49.28 | -0.319999999999993 | 13 |
| JU397 | 49.28 | -0.319999999999993 | 13 |
| JU406 | 49.28 | -0.319999999999993 | 13 |
| JU440 | 48.715 | 1.56 | 134 |
| JU561 | 48.71 | -3.81 | 39 |
| JU642 | 48.84 | 2.5 | 48 |
| JU751 | 48.84 | 2.5 | 48 |
| JU774 | 38.683 | -9.33999999999997 | 12 |
| JU775 | 38.7175 | -9.14859999999999 | 77 |
| JU778 | 38.719 | -9.14909999999998 | 65 |
| JU782 | 38.7191 | -9.15030000000002 | 85 |
| JU792 | 43.06 | 0.24 | 441 |
| JU830 | 48.52 | 9.05 | 362 |
| JU847 | 48.46 | 7.461 | 230 |
| KR314 | 49.28 | -123.13 | 45 |
| LKC34 | -18 | 46 | 687 |
| MY1 | 52.54 | 7.31 | 32 |

| | | | |
|---|---|---|---|
| MY10 | 51.96 | 7.53 | 75 |
| MY16 | 51.93 | 7.57 | 62 |
| MY18 | 51.96 | 7.53 | 75 |
| MY2212 | 54.346355 | 10.117737 | 22 |
| MY23 | 51.96 | 7.53 | 75 |
| MY2453 | 51.950777 | 7.536628 | 72 |
| MY2530 | 54.346355 | 10.117737 | 22 |
| MY2535 | 51.950723 | 7.599099 | 68 |
| MY2573 | 51.950723 | 7.599099 | 68 |
| MY2693 | 54.3491 | 10.11505 | 21 |
| MY2741 | 54.3491 | 10.11505 | 21 |
| MY679 | 54.346355 | 10.117737 | 22 |
| MY920 | 54.346355 | 10.117737 | 22 |
| NIC 1 | 43.279 | 5.3543 | 14 |
| NIC 1107 | 43.716278 | 7.266935 | 63 |
| NIC 166 | 46.72722 | 6.89775 | 684 |
| NIC 195 | 38.545874 | -28.37322 | 103 |
| NIC 199 | 37.745428 | -25.19929 | 18 |
| NIC 2 | 43.279 | 5.3543 | 14 |
| NIC 207 | 43.720505 | 7.24045 | 185 |
| NIC 231 | 43.720505 | 7.24045 | 185 |
| NIC 236 | 39.242006 | -9.31341700000002 | 14 |
| NIC 242 | 38.691765 | -9.31608999999997 | 11 |
| NIC 251 | 38.66621 | -28.151364 | 230 |
| NIC 252 | 38.661313 | -28.147949 | 213 |
| NIC 255 | 43.714834 | 7.266779 | 63 |
| NIC 256 | 38.71829 | -9.14875000000001 | 77 |
| NIC 258 | 38.71829 | -9.14875000000001 | 77 |
| NIC 259 | 38.71829 | -9.14875000000001 | 77 |
| NIC 260 | 38.71829 | -9.14875000000001 | 77 |
| NIC 261 | 38.71829 | -9.14875000000001 | 77 |

| | | | |
|---|---|---|---|
| NIC 262 | 38.71829 | -9.14875000000001 | 77 |
| NIC 265 | 38.71829 | -9.14875000000001 | 77 |
| NIC 266 | 38.69237 | -9.31592000000001 | 11 |
| NIC 267 | 38.69237 | -9.31592000000001 | 11 |
| NIC 268 | 38.69237 | -9.31592000000001 | 11 |
| NIC 269 | 38.69237 | -9.31592000000001 | 11 |
| NIC 271 | 38.69237 | -9.31592000000001 | 11 |
| NIC 272 | 39.769459 | -8.75635599999998 | 129 |
| NIC 274 | 39.769459 | -8.75635599999998 | 129 |
| NIC 275 | 39.769459 | -8.75635599999998 | 129 |
| NIC 276 | 39.769459 | -8.75635599999998 | 129 |
| NIC 277 | 43.716441 | 7.266138 | 40 |
| NIC 3 | 43.3636 | 5.3215 | 54 |
| NIC 513 | 38.776038 | -9.16713600000003 | 77 |
| NIC 514 | 38.776038 | -9.16713600000003 | 77 |
| NIC 522 | 36.535679 | -6.304441 | 0 |
| NIC 523 | 36.535679 | -6.29831000000001 | 24 |
| PS2025 | 34.19 | -118.13 | 425 |
| PX179 | 44.035 | -123.058 | 262 |
| QG2075 | 33.951 | -83.376 | 192 |
| QG536 | 37.7679 | -122.4415 | 157 |
| QG556 | 34.421629 | -119.702021 | 26 |
| QG557 | 34.421629 | -119.702021 | 26 |
| QW947 | -33.4213 | -70.6106 | 625 |
| QX1212 | 37.7502 | -122.4331 | 59 |
| QX1233 | 37.8804 | -122.2838 | 34 |
| QX1791 | 20.6344 | -156.3935 | 631 |
| QX1792 | 20.70554 | -156.35475 | 882 |
| QX1793 | 20.70559 | -156.35678 | 866 |
| QX1794 | 20.70559 | -156.35678 | 866 |
| RC301 | 47.99 | 7.84 | 268 |

| | | | |
|---|---|---|---|
| WN2001 | 51.954422 | 6.303406 | 15 |
| WN2002 | 51.975285 | 5.694834 | 29 |
| WN2033 | 51.975 | 5.694 | 29 |
| XZ1513 | 22.14787 | -159.63105 | 1230 |
| XZ1514 | 22.149 | -159.668 | 995 |
| XZ1516 | 22.149 | -159.668 | 995 |

**[Table S8] Comparison of AICc scores from analyzed MLPE models for pairwise genetic distances, with beta coefficients, *t*-statistics, and *p* values of fixed factors included in the top model.**

| Model | DF | AICc | β ± SE | *t* | *p* value |
|---|---|---|---|---|---|
| **Genetics ~ Geography** | **4** | **-117842.6** | | | |
| *Geography* | | | 0.02 ± 0.00 | 38.76 | < 0.001 |
| Genetics ~ Environment | 5 | -117834.0 | | | |
| Genetics ~ Geography + Environment | 4 | -116496.0 | | | |

**[Table S9] Comparison of AICc scores from analysed MLPE models for pairwise phenotypic distances, with beta coefficients, *t*-statistics, and *p* values of fixed factors included in the top models.**

| Model | DF | AICc | β ± SE | *t* | *p* value |
|---|---|---|---|---|---|
| **Aggregation Magnitude ~ Genetics** | **4** | **35743.20** | | | |
| *Genetics* | | | 2.12 ± 0.50 | 4.23 | < 0.001 |
| Aggregation Magnitude ~ Genetics + Elevation | 5 | 35752.17 | | | |
| Aggregation Magnitude ~ Genetics + Geography | 5 | 35749.25 | | | |
| Aggregation Magnitude ~ Genetics + Elevation + Geography | 6 | 35758.41 | | | |
| Aggregation Timescale ~ Genetics | 4 | 39913.12 | | | |
| **Aggregation Timescale ~ Genetics + Elevation** | **5** | **35532.22** | | | |
| *Genetics* | | | 1.19 ± 0.50 | 2.38 | 0.017 |
| *Elevation* | | | 0.06 ± 0.01 | 6.91 | < 0.001 |
| Aggregation Timescale ~ Genetics + Geography | 5 | 35571.36 | | | |
| Aggregation Timescale ~ Genetics + Elevation + Geography | 6 | 35537.78 | | | |
